# Functional divergence of plant expansins as revealed by genetic complementation of root-hair tip growth in *Arabidopsis thaliana*

**DOI:** 10.64898/2026.04.01.715922

**Authors:** Ke Zhou, Nathan K. Hepler, Moyan Jia, Daniel J. Cosgrove

**Affiliations:** Department of Biology, 208 Mueller Laboratory, The Pennsylvania State University, University Park, PA 16802

**Author notes:** Corresponding author: Daniel Cosgrove, Department of Biology, 208 Mueller Laboratory, The Pennsylvania State University, University Park, PA 16802.; **Email:**. Contributed equally to this work (Ke Zhou, Nathan K. Hepler). **Author Contributions:** K.Z., N.K.H and D.J.C. designed the research; K.Z., N.K.H. and M.J. performed the research; K.Z., N.K.H and D.J.C. analyzed data; K.Z., N.K.H and D.J.C. wrote the paper; D.J.C. used Nature Research Assistant Beta for shortening the abstract and identifying unclear technical details in the results section. D.J.C. supervised the research and acquired funding.

**Keywords:** expansin, cell wall, root hair growth, *Arabidopsis thaliana*, genetic complementation

## Abstract

Plant cell wall expansion is essential for plant growth, with expansins promoting wall loosening and creep during diffuse growth. Expansin’s role in tip growth, such as root hair elongation, is less clear. Here we used CRISPR/Cas9 to knock out root-hair specific α-expansins EXPA7 and EXPA18 in *Arabidopsis thaliana*, abolishing root hair elongation. Complementation with expansin genes from various clades (driven by the *EXPA7* promoter) revealed functional differences: some fully restored root hair growth, others only partially restored growth, while others failed, notably two EXPA clades [clade-VIII (EXPA13) and clade-IX (EXPA20)] lacking a conserved Asp considered essential for expansin-induced wall enlargement. Phylogenetic analysis indicated loss of this Asp predated angiosperms. Mutation of this Asp in EXPA7 confirmed its necessity for wall loosening. Members of the other expansin families (EXPB, EXLA and EXLB) failed to restore root hair growth. Chimeric fusions of EXPAs with mCherry revealed differences in trafficking patterns and wall binding among EXPA clades. These results demonstrate functional differences among expansin families and among EXPA clades and demonstrate a genetic platform to analyze functionalities (trafficking, wall binding, wall extension) of EXPA and other wall-modifying proteins during root-hair tip growth.

**Significance Statement:** Plant cells can enlarge >100-fold during growth, a process controlled by proteins called expansins. Plants have many expansin proteins that are difficult to study in isolation and whose specific activities and properties have remained elusive. Here, we engineered a root-hairless *Arabidopsis* mutant as a genetic platform to assess expansin activity *in vivo*. By introducing various expansins with a fluorescent tag into this mutant, we uncovered an unexpected functional diversity across the expansin superfamily. While some expansins successfully restored root hair growth, others failed, potentially due to differences in protein trafficking, cell wall binding, or evolutionary loss of critical residues required for wall loosening. These findings point to a divergence of expansin functions during >140 million years of plant evolution.

## Introduction

Plant cell growth involves cell wall relaxation and yielding to turgor pressure, driving cell water uptake and irreversible wall enlargement (1-3). This process enables plant cells to enlarge >100-fold in volume, critical for tall stems, deep roots and large leaf canopies. Growing cell walls are both strong and extensible – properties enabled by a cohesive network of cellulose microfibrils embedded in a hydrated matrix of hemicelluloses and pectins (1-4). Two distinctive patterns of cell enlargement are commonly identified as (a) *diffuse growth*, where the entire wall surface enlarges - exemplified by epidermal cells in elongating hypocotyls - and (b) *tip growth* - exemplified by elongating root hairs - where expansion is largely restricted to the tip of tubular cells (5, 6). Recent studies have emphasized the local secretion, incorporation and remodeling of pectins in tip growth (7-9) whereas expansin-mediated enlargement of the cellulose network is emphasized for diffuse growth (2, 10-12). In this study we examine the requirement for expansins in tip growth of root hairs. We develop a complementation assay for expansin function based on root hair growth of a hairless mutant and use this platform to assess biological functionalities of diverse members of the expansin superfamily in *Arabidopsis thaliana*.

Expansins were discovered as protein mediators of ‘acid growth’, pH-dependent extension of plant cell walls commonly linked to auxin-induced diffuse cell growth (12-14). Ubiquitous in land plants, expansins comprise four multi-gene families: α-expansin (EXPA), β-expansin (EXPB), expansin-like A (EXLA) and expansin-like B (EXLB) (15-18). Native EXPA proteins from diverse species can facilitate acid-induced wall extension *in vitro*, and this is widely considered a defining characteristic of EXPAs, whereas the actions of EXPB, EXLA and EXLB proteins are less clear (10); EXPA proteins are encoded by 26 genes in *Arabidopsis thaliana* and 34 in *Oryza sativa* (rice) (17, 19). This gene multiplicity raises long-unanswered questions: Do different EXPAs possess functionally distinct activities, perhaps tuned to different wall polysaccharides? Or do EXPAs possess equivalent activities, fulfilling different biological roles by virtue of different patterns of expression?

The first possibility is suggested by the deeply conserved EXPA phylogeny, encompassing twelve monophyletic clades (denoted EXPA-I to EXPA-XII) found throughout angiosperms (20-23). Biochemical specialization of expansins at the family level is supported by the fact that EXPAs bind and loosen cellulose networks (24, 25) whereas maize EXPB1 loosens arabinoxylans in grass cell walls (26, 27). Distinctive expression patterns for different expansin genes support the second possibility (11, 17, 28-31).

Progress on this topic has been limited by recalcitrance of plant expansins to recombinant expression of active protein in heterologous expression systems (10), largely restricting *in-vitro* analyses to native expansin proteins that can be isolated in sufficient quantity and purity for such assessments. Because plant expansins are generally present in miniscule amounts and bind tightly to cell walls (14, 32), this approach is narrowly circumscribed. *In-silico* modeling of EXPA binding to polysaccharides has been reported (33), but lacks experimental validation. The discovery of microbial expansins (34) which are readily expressed heterologously (35, 36), enabled detailed structure-function analyses of these proteins by mutagenesis combined with *in-vitro* wall extension assays (37). However, microbial expansins studied to date exhibit low activity and low sequence similarity to plant expansins (15, 38). Hence, studies of microbial expansins cannot answer questions about the unique wall-loosening activities and biological functions of EXPAs.

To develop an alternative approach based on genetics, we created a root-hairless Arabidopsis line by stacking null mutations of two clade EXPA-X genes that are specifically expressed in root hairs, enabling assessment of expansin activity by genetic complementation of the hairless phenotype. With EXPA-mCherry fusions (28) we also monitored EXPA subcellular trafficking, polarized secretion and wall-binding properties in root-hairs, revealing an unexpected diversity in EXPA functionalities.

## Results

### Root hair elongation in Arabidopsis requires expression of an EXPA

Previous results showed that the two genes from clade EXPA-X are expressed exclusively in trichoblasts (root hair progenitors) (30, 39) and that RNAi suppression reduced root hair length by 25-50% (40). To fully knock out the two EXPA-X genes of *Arabidopsis thaliana* Col-0 (30), we employed CRISPR/Cas9-mediated genome editing using three polycistronic tRNA-gRNA scaffolds (41) to target *EXPA7, EXPA18* or both genes (see **Fig. S1A**,**B** and **Methods** in **Supplementary Materials**). Homozygous lines with disrupted genes *expa7-1, expa7-2, expa18-1*, and *expa7-3/expa18-3* (**Fig. S1C**,**D**) were isolated and characterized. Under our optimized growing conditions (see Methods), root hairs of the single-gene mutants *expa7-1, expa7-2* and *expa18-1* grew to the same length as Col-0, ∼550 μm (**Fig. 1A,B**), whereas under suboptimal growth conditions, the single-gene knockout mutants had shorter root hairs than Col-0 (**Fig. S2**), a pattern consistent with previous EXPA-X knockdown studies (40, 42). EXPA dosage effects may be sensitive to growth conditions. In two different double mutants (*expa7-3/expa18-3* and *expa7-2*/*expa18-1)*, root hairs initiated at the same location and density as in Col-0 (**Fig. S3A-D**), but failed to elongate (**Fig. 1A,B**). From these results we conclude that root-hair tip growth in Arabidopsis requires expression of at least one EXPA.

**Figure 1.**
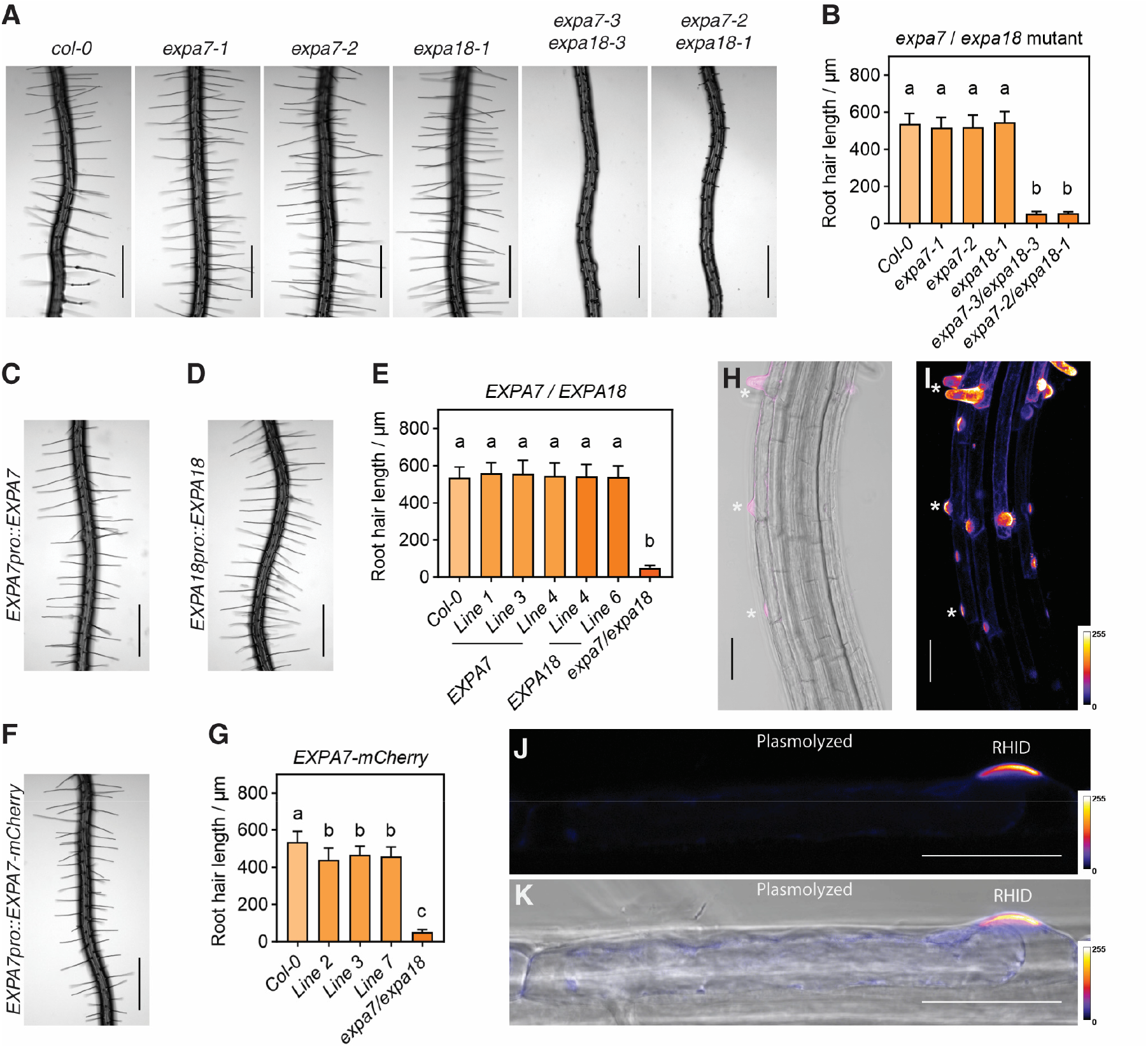
EXPA7 and EXPA18 requirement for root hair growth and subcellular localization of EXPA7. (**A** and **B)** Root hairs (A) and root hair length (B) of Col-0 (n=43), *expa7-1* (n=33), *expa7-2* (n=49), *expa18-1* (n=41), *expa7-3/expa18-3* (n=37), and *expa7-2/expa18-1* (n=48). (**C** to **E**) Restoration of root hair growth in *expa7/expa18* mutant by *EXPA7pro::EXPA7* (n=30, 46, 37) and *EXPA18pro::EXPA18* (n=44, 45). (**F** to **G**) Restoration of root hair growth in *expa7/expa18* mutant by *EXPA7pro::EXPA7-mCherry* (n=34, 25, 21). **(H and I)** EXPA7-mCherry localization in the root region spanning the transition from the elongation zone: single optical section (H) and maximum-intensity projection (I). Asterisks mark EXPA7-mCherry accumulation at different stages of root hair development. **(J and K)** EXPA7-mCherry localization in a bulging trichoblast plasmolyzed with 500 mM mannitol (J, fluorescence channel; K, merged channel). Fluorescence signals in (I to K) are displayed using the Fire LUT (ImageJ), a pseudocolor scale in which pixel intensities are represented from low to high relative fluorescence signal, with darker colors indicating lower signal and warmer colors indicating higher signal; color bars indicate the displayed intensity range from 0 to 255. Asterisks mark cell wall–associated EXPA7-mCherry at the RHID. Numerical data are means ± SD. One-way ANOVA followed by Tukey’s HSD multiple comparison test was used; different letters denote statistically significant differences (P < 0.0001) in (B), (E), and (G). Scale bars: (A, C, D, F), 500 μm; (H, I, J, K), 50 μm.

When *EXPA7* or *EXPA18* genes, driven by their native promoters (30), were introduced into the *expa7-3/expa18-3* line (henceforth designated *expa7/expa18* for brevity), root hair elongation was restored (**Fig. 1C-E**), confirming that *EXPA7* and *EXPA18* are redundantly required for root hair elongation beyond the initiation phase. To assess the evolutionary conservation of clade EXPA-X function, we tested *SmEXPA5* - an ortholog of *AtEXPA7* (22) from the lycophyte *Selaginella moellendorffii*, a primitive vascular plant. When driven by *EXPA7pro* (30), *SmEXPA5* complemented the root hairless phenotype of *expa7/expa18* (**Fig. S4**), demonstrating the ancient functional similarity of clade EXPA-X proteins (42), predating angiosperm evolution. These results extend previous work implicating EXPAs in root-hair growth (30, 40, 42, 43).

### EXPA7 is targeted to the cell wall of the root hair initiation domain (RHID) and the root-hair tip

To visualize the spatial pattern of EXPA trafficking and localization, we fused EXPA7 with mCherry using a short linker (encoding GGSGGGSGG). When driven by *EXPA7pro* (30) and introduced into *expa7/expa18, EXPA7-mCherry* largely restored root hair elongation (**Fig. 1F,G**). Fluorescence initially appeared precisely at the site of root hair initiation preceding and coincident with wall bulging and subsequently increased in intensity with elongation of the nascent root hair (**Fig. 1H,I**) and was localized to the tip for later stages of elongation (**Fig. S5**). Plasmolysis of cells with emerging bulges showed that the fluorescence was stably bound to the cell wall of the root hair (**Fig. 1J,K**). These results indicate that EXPA7-mCherry is functional and stably binds to the cell wall in a spatiotemporal pattern consistent with a role in facilitating wall extension during root hair outgrowth and subsequent elongation.

### EXPAs in Clades I, II, and IV are functionally equivalent to EXPA7 for root hair growth

The twenty-six EXPAs in Arabidopsis are members of phylogenetic clades EXPA-I to EXPA-XII (**Table 1**) (23). Whether proteins from different EXPA clades possess equivalent subcellular trafficking, binding and wall growth activities is unknown. We addressed this question by complementation of *expa7/expa18* using *EXPA7pro::EXPAx-mCherry* constructs; root hair phenotypes and mCherry fluorescence patterns were assessed in homozygous lines. If EXPAs from different clades possess equivalent activities, we would expect complementation as found for EXPA7-mCherry (**Fig. 1F-H**). This proved to be the case for EXPAs from some but not all clades.

**Table 1.** Family and cladal organization of expansins in *Arabidopsis thaliana* and summary of root hair growth complementation results.

|  |  |  |
| --- | --- | --- |
| EXPA | Clade I | <u>EXPA1</u><sup>+</sup>, EXPA5, <u>EXPA10</u><sup>+</sup>, EXPA15 |
|  | Clade II | <u>EXPA14</u><sup>+</sup> |
|  | Clade III | <u>EXPA2</u><sup>1</sup>, EXPA8 |
|  | Clade IV | EXPA3, <u>EXPA4</u><sup>+</sup>, EXPA6, EXPA9, EXPA16 |
|  | Clade V | <u>EXPA11</u><sup>1</sup> |
|  | Clade VI | EXPA17 |
|  | Clade VII | <u>EXPA12</u><sup>1</sup> |
|  | Clade VIII | <u>EXPA20</u><sup>0</sup> |
|  | Clade IX | <u>EXPA13</u><sup>0</sup> |
|  | Clade X | <u>EXPA7</u><sup>+</sup>, <u>EXPA18</u><sup>+</sup> |
|  | Clade XII | EXPA19<sup>ps</sup>, <u>EXPA21</u><sup>0</sup>, EXPA22, EXPA23, <u>EXPA24</u><sup>0</sup>, EXPA25, EXPA26 |
| EXPB | Clade I | EXPB2, EXPB4, EXPB5, EXPB6 |
|  | Clade II | <u>EXPB1</u><sup>0</sup>, EXPB3 |
| EXLA | Clade I | <u>EXLA1</u><sup>0</sup>, EXLA2, EXLA3 |
| EXLB | Clade I | <u>EXLB1</u><sup>0</sup> |

Notes: EXPA17 is phylogenetically close to EXPA12, so was not selected for study; EXPA19<sup>ps</sup> is a pseudogene in ecotype Col-0 but not Landsberg erecta. Clade XI is absent in Arabidopsis.

Chimeric mCherry fusions with EXPA1 (Clade-I), EXPA14 (Clade-II), and EXPA4 (Clade-IV) were successfully expressed (**Fig. S6A**) and restored root hair elongation of the *expa7/expa18* line as observed for EXPA7-mCherry (**Fig. 2A-C; 2G**). As with EXPA7-mCherry, fluorescence accumulated specifically at the RHID and was stably bound to the cell wall in each of these cases (**Fig. 2A-C**). We also tested EXPA10 from Clade-I, obtaining results similar to its paralog EXPA1 (**Fig. S7A-E**). These results demonstrate the functional equivalence of six EXPAs from four clades (I, II, IV and X) in terms of protein targeting, cell wall binding and promoting root hair elongation.

**Figure 2.**
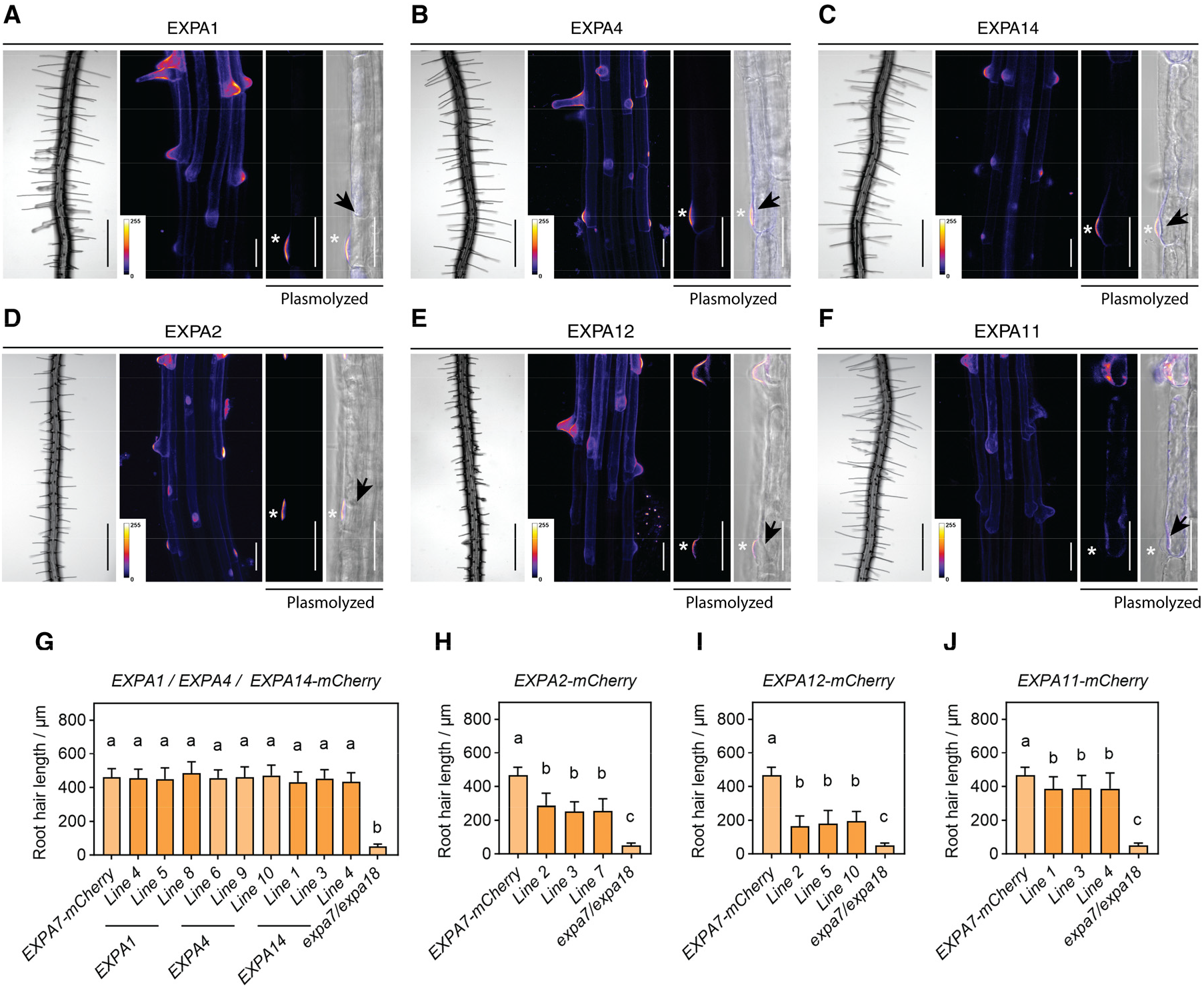
Some EXPAs rescue root hair growth of *expa7/expa18* mutant. (**A** to **F**) Restoration of root hair growth and subcellular localization of mCherry-fused EXPA1 (A), EXPA4 (B), EXPA14 (C), EXPA2 (D), EXPA12 (E), and EXPA11 (F) in *expa7/expa18*. In panels A to F, from left to right: root hairs; a maximum-intensity confocal projection of the root region spanning the transition from the elongation zone to the differentiation zone; the fluorescence channel (EXPA1, EXPA4, EXPA14, EXPA2, EXPA12) or bright-field channel (EXPA11); and merged fluorescence and bright-field channels of a plasmolyzed bulging trichoblast. Fluorescence signals are displayed using the Fire LUT (ImageJ). Asterisks indicate the plasmolyzed cell wall at the RHID. Scale bars: 500 μm, 50 μm, 50 μm, and 50 μm (from left to right). (**G**) Root hair length of independent lines expressing mCherry-fused EXPA1 (n=50, 28, 48), EXPA4 (n=23, 20, 24), and EXPA14 (n=31,18, 20); (**H**) EXPA2 (n=41, 30, 27); (**I**) EXPA12 (n=23, 21, 34); and (**J**) EXPA11 (n=50, 42, 41) in *expa7/expa18* mutant. EXPA7-mCherry line 2 (n=31) serves as positive control. For statistical analysis, data are presented as means ±SD with One-way ANOVA followed by Dunnett’s **(G)** or Tukey’s (H, I, J) HSD multiple comparison tests; different letters denote statistically significant differences (P<0.0001).

### EXPAs from Clades III, V and VII differ functionally from EXPA7

In contrast to the above results, EXPA2-mCherry (Clade-III) lines, which successfully expressed intact mCherry-fusion protein (**Fig. S6B**), only partially rescued root hair growth of *expa7/expa18* (**Fig. 2D,H**), yet fluorescence accumulated at the RHID and was bound to the cell wall in the manner of EXPA7-mCherry (**Fig. 2D**). Time course analysis showed that EXPA2-mCherry root hairs elongated more slowly than EXPA7-mCherry controls (**Fig. S8**). Similar results were obtained with EXPA12-mCherry (Clade-VII), which showed even weaker rescue of root hair growth. (**Fig. 2E,I, Fig. S6B**). The basis for the reduced growth activity of EXPA2-mCherry and EXPA12-mCherry is unknown; it might be due to differences in intrinsic protein activity, secretion efficiency or binding properties within the root hair wall. Western blots indicate it is not due to less protein (**Fig. S6B**). A different pattern was observed for EXPA11-mCherry (Clade-V), which largely complemented the growth phenotype of *expa7/expa18* yet displayed only weak accumulation and binding at the RHID (**Fig. 2F,J, Fig. S6B**). Plasmolysis of cells in the mid-differentiation zone of the root showed appreciable accumulation of fluorescence between the cell wall and the retracted plasma membrane (**Fig. S9**), indicating EXPA11-mCherry secretion but reduced wall binding. Nevertheless, root hair elongation was only slightly diminished compared with EXPA7-mCherry (**Fig. 2J**), suggesting that strong wall binding is not essential for EXPA action in this assay or that small amounts of mobile EXPA suffice for tip growth.

These results show that trafficking, wall-binding and wall-extension activities vary appreciably among EXPAs, revealing an unanticipated functional diversity. Even more extreme differences were found with Clades VIII and IX.

### EXPAs from Clades VIII and IX fail to restore root hair growth and lack a highly conserved Asp

EXPA13 (Clade IX) and EXPA20 (Clade VIII) completely failed to rescue root hair growth in *expa7/expa18* (**Fig. 3A**,**B**; quantitation in **Fig. S10A**,**B**). These two clades are closely related phylogenetically (15, 23). Fluorescence was not localized to the RHID and bound only weakly to the cell wall (**Fig. 3A,B**). Much of the mCherry fluorescence appeared to be cytoplasmic and observations of plasmolyzed cells in the mid-differentiation zone showed intracellular fluorescence with only weak binding to the cell wall of bulges, which failed to elongate beyond the initiation stage (**Fig. 3A,B**). Western blot results showed that an appreciable portion of EXPA13-mCherry and EXPA20-mCherry was proteolytically cleaved (**Fig. S6C**), potentially a result of intracellular protein digestion due to poor secretion (Fig. 3A, B). These results indicate that EXPA13 and EXPA20 differ markedly from EXPA7 in terms of protein trafficking, secretion, wall binding and wall enlargement activity in Arabidopsis root hairs.

**Figure 3.**
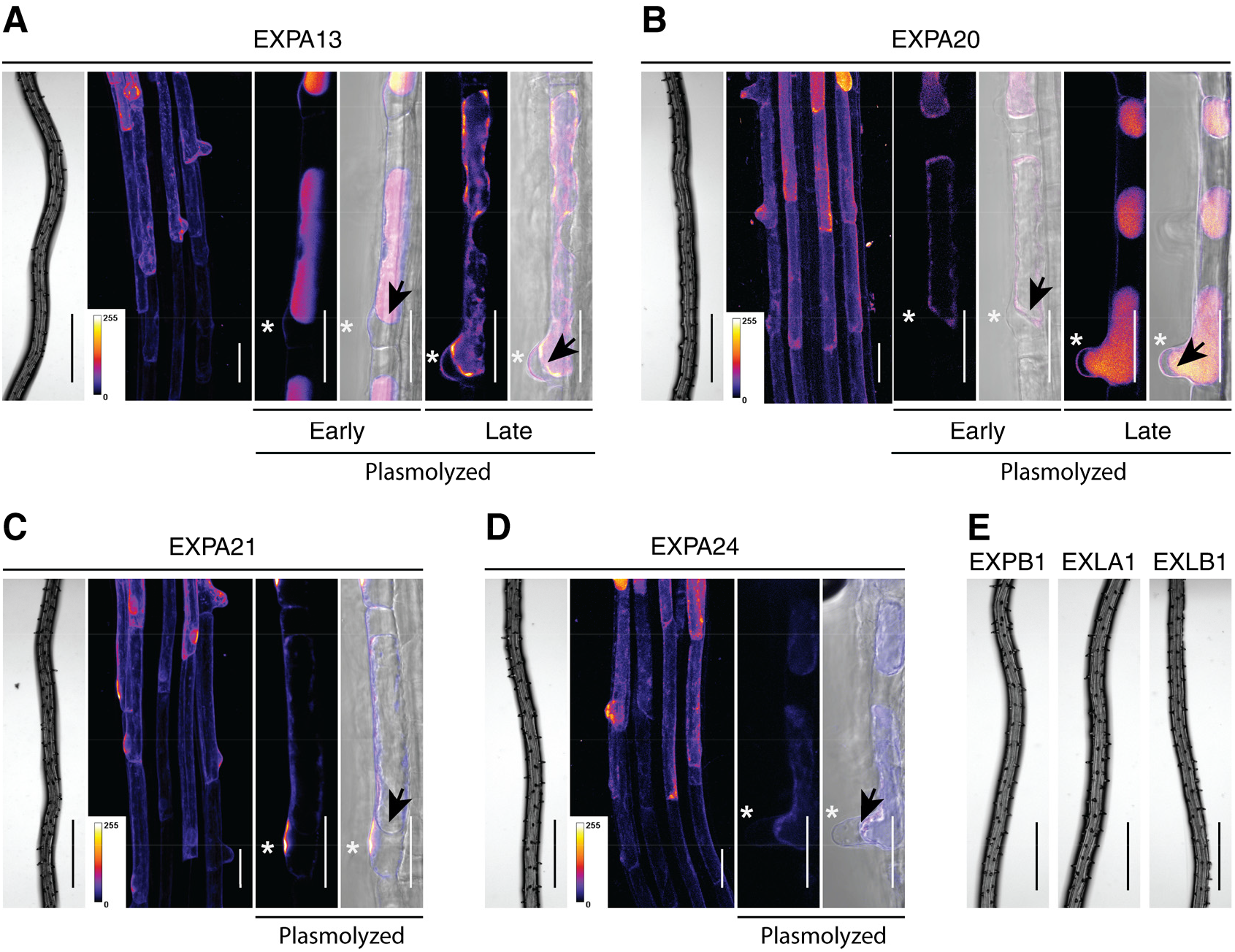
Some expansin fail to rescue root hair growth of *expa7/expa18*. **(A to D)** Root hair growth of *expa7/expa18* expressing mCherry-fused EXPA13 (A), EXPA20 (B), EXPA21 (C), or EXPA24 (D), and subcellular localization of the fusion proteins. Panels A–D from left to right show root hairs; a maximum-intensity confocal projection of the root region spanning the transition from the elongation zone to the differentiation zone; the fluorescence channel (EXPA21, EXPA24) or bright-field channel (EXPA13, EXPA20); and merged fluorescence and bright-field channels of a plasmolyzed trichoblast. For EXPA13 (A) and EXPA20 (B), additional panels show plasmolyzed trichoblasts at two root hair developmental stages: Early, corresponding to newly bulging trichoblasts in the root hair initiation region within the elongation-to-differentiation transition region; and Late, corresponding to trichoblasts in the mature differentiation zone, where root hairs failed to undergo normal elongation and remained as bulges. Fluorescence signals are displayed using the Fire LUT (ImageJ). Asterisks indicate the plasmolyzed cell wall at the RHID. (**E**) Lack of root hair growth of *expa7/expa18* seedlings expressing *EXPB1, EXLA1*, or *EXLB1*. Scale bars, 500 μm (root hair overviews in A to E); 50 μm (all other panels).

Previously, mutagenesis of the microbial expansin BsEXLX1 from *Bacillus subtilis* showed that a highly conserved Asp residue (D82) is essential for its wall loosening action, measured as induction of cell wall creep (37). D82 is located in a presumed active center of expansin domain-1 which is proposed to facilitate twisting of glucan chains as a part of its wall-loosening action (10, 44). Structurally, D82 corresponds to the highly conserved Asp in the ‘HFD’ motif characteristic of EXPA and EXPB (**Fig. 4A**, in green box) (15, 26, 34) and corresponds to the catalytic Asp in family-45 glycosyl hydrolases (15, 45). In EXPA13 this Asp is replaced by valine (V) and in EXPA20 by glutamic acid (E) (**Fig. 4A**).

**Figure 4.**
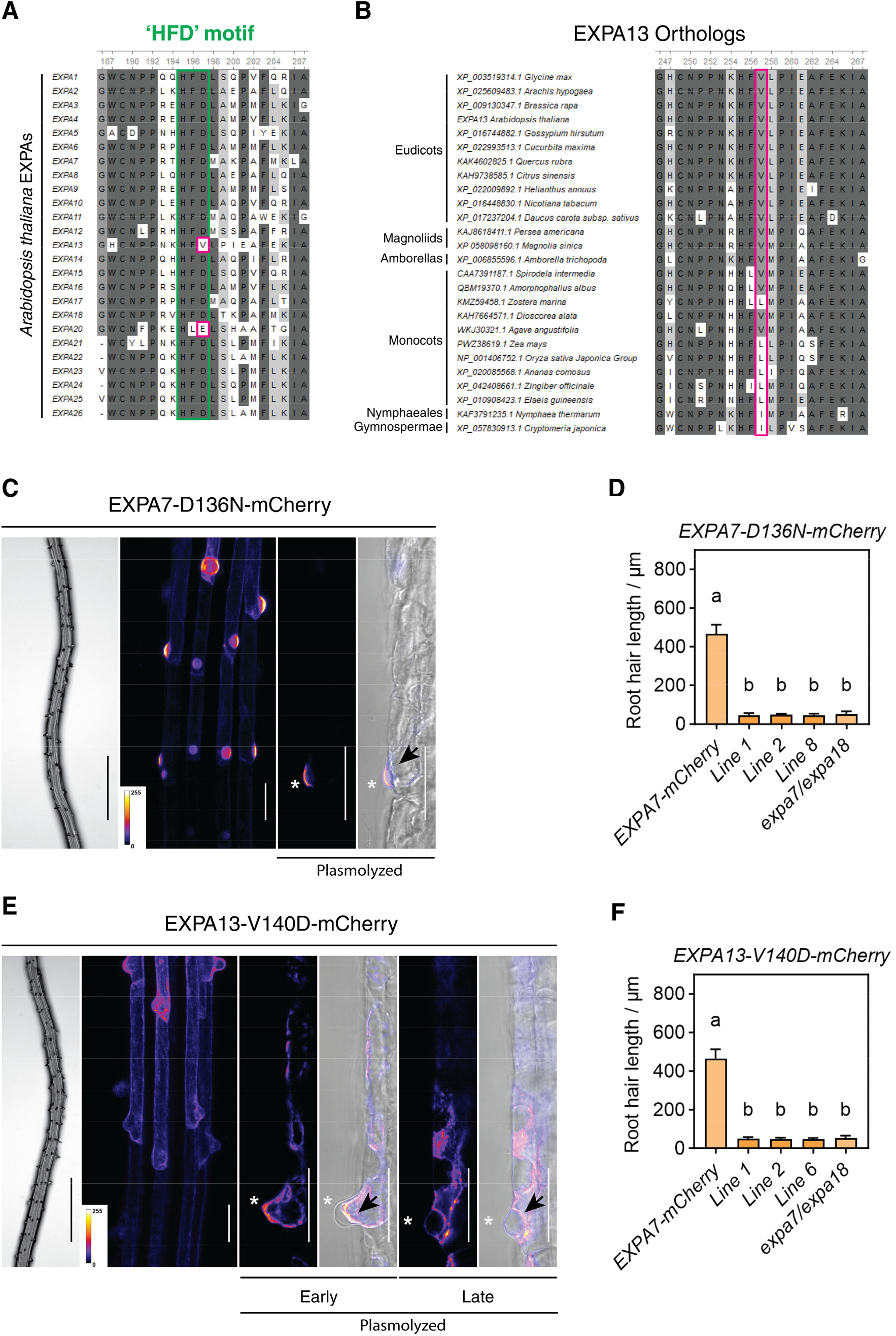
Loss of critical Asp residue in EXPA13 and EXPA20. (**A**) Partial sequence alignment of Arabidopsis EXPAs. The conserved ‘HFD’ motif is boxed in green; and the divergent residues in EXPA13 and EXPA20 are boxed in purple. (**B**) Amino-acid sequence alignment of EXPA13 orthologs from angiosperms and basal land plants. Conserved Val residues are boxed in purple. (**C** and **E**) Root hair growth of *expa7/expa18* expressing EXPA7-D136N-mCherry (C) or EXPA13-V140D-mCherry (E), and subcellular localization of the fusion proteins. In **(C, E)**, panels from left to right: root hairs; a maximum-intensity confocal projection of the root region spanning the transition from the elongation zone to the differentiation zone; the fluorescence channel **(C)** or bright-field channel **(E)**; and merged fluorescence and bright-field images of plasmolyzed trichoblasts. For EXPA13-V140D **(E)**, additional panels show plasmolyzed trichoblasts at two root hair developmental positions: Early, corresponding to newly bulging trichoblasts in the root hair initiation region within the elongation-to-differentiation transition region; and Late, corresponding to trichoblasts in the mature differentiation zone, where root hairs in these non-rescuing lines failed to undergo normal elongation and remained as bulges. Fluorescence signals are displayed using the Fire LUT (ImageJ). Scale bars, 500 μm (root hair overviews in C and E); 50 μm (all other panels). (**D** and **F**) Root hair length of *expa7*/expa18 expressing EXPA7-D136N-mCherry (D; n=32, 29, 22) or EXPA13-V140D-mCherry (F; n=22, 26, 19) ; EXPA7-mCherry line 2 (n=31) serves as positive control. For statistical analysis, data are means ± SD. One-way ANOVA was performed, followed by Tukey’s HSD multiple comparison tests. Different letters indicate significantly different groups (P<0.0001).

Bioinformatic analysis showed that this Asp is consistently absent in putative orthologs of EXPA13 and EXPA20 throughout angiosperms, from basal angiosperms (Amborellales, Nymphaeales) to monocots and eudicots (**Fig. 4B** and **Fig. S11**). Phylogenetic analyses confirmed these sequence variations as consistent characteristics of Clades VIII and IX (**Fig. S12**) which are sister clades. We also identified EXPA13 and EXPA20 orthologs from the gymnosperm *Cryptomeria japonica*, indicating these conserved sequence variations predate angiosperm diversification.

In EXPA7 we mutated the homologous Asp (D136) to Asn (N) and introduced *EXPA7pro::EXPA7*-*D136N-mCherry* into *expa7/expa18*. Fluorescence accumulated at the RHID and was bound to the cell wall as with EXPA7-mCherry, but the D136N protein failed to restore root hair growth (**Fig. 4C,D, Fig. S6D**), demonstrating that the Asp in the conserved HFD motif is required for wall loosening activity of EXPA7 *in vivo*.

We also carried out a reciprocal test with EXPA13, mutating the corresponding Val residue to Asp, creating *EXPA7pro::EXPA13-V140D-mCherry* and introducing it into *expa7/expa18*. This did not restore root hair elongation and fluorescence was not targeted to the RHID, but the protein instead underwent substantial degradation (**Fig. 4E,F, Fig. S6D**), similar to the unmodified EXPA13, perhaps as a consequence of poor secretion. Evidently the absence of this Asp residue in EXPA13 is not the sole basis for its lack of wall activity in this assay.

### EXPA21 and EXPA24 (Clade XII) also fail to restore root hair growth

*EXPA21*-*EXPA26* genes are tandem duplications on chromosome V (23). We introduced *EXPA21-mCherry* and *EXPA24-mCherry* into *expa7/expa18* under control of the *EXPA7* promoter, but neither construct restored root hair growth (**Fig. 3C,D** and **Fig. S6C, Fig. S10C**,**D**).

Microscopy showed that EXPA21-mCherry accumulated preferentially at the RHID and bound to the cell wall (**Fig. 3C**).Notably, western blot analysis revealed multiple EXPA21-mCherry bands, which may reflect differential processing or post-translational modification of the fusion protein. Because several members of the expansin superfamily were identified in previous N-glycoproteomic datasets (46), glycosylation represents one possible explanation, but this was not further studied here. EXPA24-mCherry did not accumulate at the RHID wall (**Fig. 3D**), perhaps because of weak wall binding, degradation (**Fig. S6C**) or inefficient secretion. Further analysis of the EXPA24 sequence revealed a low-complexity repetitive region of ∼60 residues immediately following the predicted signal peptide (**Fig. S13**), which might affect trafficking and secretion.

These results hint at functional differences within this group of tandem genes.

### EXPB, EXLA and EXLB genes do not restore root hair elongation

Little is known about wall-related activities of these expansin families in Arabidopsis (10) except for a recent report that recombinant AtEXLA1 may have very weak creep activity in vitro (47); its relevance to growth in-vivo is unclear. Similarly, overexpression of AtEXLA2 with a strong constitutive promoter in Arabidopsis had little effect on cell wall creep (48). To test for wall extension activity, we introduced representative genes (*EXPB1, EXLA1, EXLB1)* from each family into *expa7/expa18* (see **Table 1**). None rescued root hair growth in these trials (**Fig. 3E** and **Fig. S14**). These results indicate functionalities distinct from EXPA7, but what those are remain an open question.

## Discussion

Root hairs elongate primarily by tip growth, a highly polarized form of localized wall expansion involving coordination of localized ion currents, calcium signaling, membrane recycling, cytoskeletal dynamics, vesicular trafficking and deposition of wall materials to the cell wall at the root-hair tip, coordination of water uptake with wall expansion, signaling by reactive oxygen species, and other sensory feedback systems [reviewed in (49, 50)]. Mechanistically, the process ultimately reduces to local wall relaxation and turgor-driven expansion of the cell wall, where stress anisotropy and mechanical anisotropy influence the pattern of localized expansion (51, 52).

Tip growth is often contrasted with diffuse growth, where entire cell wall facets enlarge in surface area (32), mediated at least in part by EXPAs (10). Arabidopsis EXPAs have previously been implicated in root hair growth by expression analysis (30) and by reduced root-hair length when EXPA expression is reduced (40, 42). The lack of root hair elongation in the *expa7/expa18* mutant indicates that the highly polarized mode of cell wall expansion in Arabidopsis root hairs requires EXPA action. This is consistent with the hypothesis that similar physical mechanisms of wall enlargement may underlie both diffuse growth and tip growth (53). Ring-like patterns of condensed pectin have recently been observed in growing root hairs (8). How pectins might influence EXPA-mediated wall expansion during tip growth is an interesting issue yet to be explored (6).

Because of technical limitations in expressing plant expansins in heterologous expression systems, detailed knowledge of expansin’s actions on cell walls has been limited to just a few members of this superfamily (10). Complementation of *expa7/expa18* circumvents this bottleneck, enabling a genetic approach to assess expansins for wall growth and for patterns of subcellular trafficking, polarized secretion and wall binding. Full or partial growth complementation was observed for proteins from seven EXPA clades, whereas proteins from three EXPA clades failed to complement (Table 1) as did members of the more divergent EXPB, EXLA and EXLB families (Table 1). Previous sequence analysis showed that EXLA and EXLB lack the highly conserved Asp in the HDF motif found in EXPA and EXPB (15), which is consistent with the lack of complementation found here (Fig. 3E). Biochemical studies showed that xylans are the preferred targets of EXPB protein (27, 54), and xylans (55) may not contribute to Arabidopsis root-hair growth. These results support the case for substantial functional differences both among expansin families and among clades within the EXPA family.

Particularly striking are results showing the lack of *expa7/expa18* complementation by EXPA13 and EXPA20 and the absence of the highly conserved Asp in the HFD motif in these two proteins and in their orthologs throughout clades EXPA-VIII and EXPA-IX. These two clades predate the origin of angiosperms. Mutation of this conserved Asp in EXPA7 rendered the protein incapable of restoring root hair growth in *expa7/expa18*, similar to results obtained for the bacterial expansin from *Bacillus subtilis* (37). This leads us to propose that EXPAs in these two ancient EXPA clades – and possibly Clade XII as well – systematically lack the classical wall-loosening activity of canonical EXPAs. These noncanonical EXPAs may possess other adaptive functions maintained by selection throughout ∼140 million years of angiosperm diversification (12, 17, 56).

## Materials and Methods

### Plant materials and growth conditions

For root hair observation, seeds were surface-sterilized and sown on half-strength Murashige and Skoog (½ MS) agar plates (pH 5.8; 0.5 g L^−1^ MES adjusted with 1 M KOH) supplemented with 1% (w/v) sucrose and 0.5% (w/v) Gelzan (G-1910, Sigma). After cold stratification at 4 °C in darkness for 2 d, plates were placed vertically and seedlings were grown under white light (100 µmol m^−2^ s^−1^) with a 16 h/8 h light/dark photoperiod at 22 °C for 4 d before imaging. Root hairs were plasmolyzed with 500 mM mannitol.

### Knocking out *EXPA7* and *EXPA18* by CRISPR/Cas9

Cloning and assembly of gRNA complexes were performed using the plasmids pGTR and pHEE401E. pGTR was a gift from Yinong Yang (Addgene plasmid # 63143 ; http://n2t.net/addgene:63143 ; RRID:Addgene_63143) and pHEE401E was a gift from Qi-Jun Chen (Addgene plasmid # 71287 ; http://n2t.net/addgene:71287 ; RRID:Addgene_71287). gRNAs targeting *EXPA7* and *EXPA18* were designed using CRISPR-PLANT (41). Two gRNAs for each were selected, both of which have a low off-targeting potential and target exons in order to increase the likelihood of frameshift or nonsense mutations. Polycistronic tRNA-gRNAs (PTGs) were created as described (41) with some modifications. Tandem tRNA-gRNAs were assembled from PCR products amplified using the tRNA and gRNA scaffold-containing plasmid pGTR. Both the PTGs and pHEE401E were digested using *BsaI*, removing pHEE401E’s 1,221-bp sgRNA region(57). The linearized vector was dephosphorylated using shrimp alkaline phosphatase and ligated alongside each PTG using T4 DNA ligase (New England Biolabs, Ipswich, MA, USA). pHEE401E-PTG ligations were recovered through the transformation of 5-alpha chemically competent *E. coli* (New England Biolabs, Ipswich, MA, USA) and selection on LB agar containing 50 mg/ml kanamycin. Plasmid was purified (Plasmid Mini kit, QIAGEN), transformed into *Agrobacterium tumefaciens* strain GV3101, and then introduced into *Arabidopsis* via the floral dip method (58, 59). The generated *Arabidopsis* lines were purified and confirmed by sequencing desired genes.

### Native gene complementation

For native gene complementation constructs, *EXPA7* (1,881-bp) and *EXPA18* (1,942-bp), including the minimal promoter and 3’-UTR regions, were cloned from genomic DNA, as previously elucidated (30). The cloned fragments were introduced into a pCAMBIA binary vector, then into *Agrobacterium tumefaciens* strain GV3101 and, consequently, the expa7-3/expa18-3 root hairless mutant via the floral dip method. Transformants were selected through supplementary hygromycin in the medium, and the introduction of native genes was confirmed by qPCR.

### Cloning expansin genes and fusion with fluorescence protein gene

To clone the selected *EXPA, EXPB, EXLA*, and *EXLB* family genes, total RNA was extracted (RNeasy Plant Mini kit, QIAGEN) from *Arabidopsis* seedlings, leaves, roots, and flowers, respectively, and then, coding sequences of desired genes were cloned from the synthesized cDNA. Genomic *EXPA21* was used instead of cDNA which did not amplify from various tissues. To drive expression in the trichoblast, a 538-bp segment upstream of the *EXPA7 coding region* was used as the trichoblast-specific promoter, as previously described (30). For visualization, we used mCherry (Accession: AHW57114.1) as the fluorescence reporter in this study. The *mCherry* gene sequence was optimized for plants and synthesized. A linker encoding a flexible polypeptide, GGSGGGSGGS, was added at its N-terminus for better fusion. Coding sequences of desired *EXPA* genes were inserted between the promoter and the optimized *mCherry* gene (*EXPA7pro::EXPAx-mCherry*) in the pCAMBIA binary vector, then introduced into *Agrobacterium tumefaciens* strain *GV3101* and subsequently into the *expa7-3/expa18* root hairless mutant via the floral dip method. Transformants were selected through supplementary hygromycin in the medium. At least ten individual lines of each transformant, screened by hygromycin resistance and fluorescence observation, were obtained and purified, and representative homozygous from three individual lines were utilized for root hair length quantitative analysis.

The coding sequence of *Selaginella moellendorffii SmEXPA5* (NCBI ACCESSION: XP_002961058) was synthesized, fused with the EXPA7 promoter (30) and transformed into the *expa7/expa18* line.

### Imaging

Seedling roots used for root hair length analysis were imaged under Olympus MVX10 Macro Zoom Microscope. For subcellular localization analysis, the seedling roots with mCherry fluorescence-labeled EXPAs were imaged by Zeiss LSM780 Axio Observer laser scanning confocal microscope (561 nm excitation and 580-652 nm emission). ZEISS ZEN 3.8 software was utilized to build the maximum intensity projection of 3D confocal stacks.

Where indicated, fluorescence images are displayed using the Fire LUT by ImageJ (1.53u version) to facilitate visualization of relative fluorescence intensity patterns. In this pseudocolor scale, pixel intensities are mapped from low to high signal, with darker colors representing lower relative fluorescence and warmer colors representing higher relative fluorescence. The accompanying color bars indicate the displayed 8-bit intensity range from 0 to 255. These pseudocolor images are more informative for visualization of fluorescence distribution compared with absolute fluorescence quantification. The color bar is autoscaled for each image and so differs between images.

### qRT-PCR of root-expressed genes

Total RNA used for qRT-PCR was extracted from four-day-old *Arabidopsis* roots using the QIAGEN RNeasy Plant Mini kit, and cDNA was synthesized using a High-Capacity cDNA reverse transcription kit (Applied Biosystems). PCR reactions were performed using PerfeCTa SYBR Green FastMix (Quantabio) on a StepONE-Plus quantitative PCR machine (Applied Biosystems) per the manufacturer’s instructions.

To amplify transcripts in seedling roots from transgenic *EXPB1, EXLA1*, and *EXLB1* constructs, we used the reverse primer localizing within the 5’-NOS terminal (Nos_qpcr_r: CGCAAGACCGGCAACAGGATT) and forward primers localizing within the desired insertion genes (EXPB1_qpcr_f: GGGACCATTCTCCGTGAAGCTC; EXLA1_qpcr_f: TTCCATCCAATTGGGAAGCTGG; EXLB1_qpcr_f: TGGATCCAATCGCCAAACGC) to assess the insertion and expression of these genes in transformants. ACT2 was utilized as the reference (ACT2_qpcr_F: CGTACAACCGGTATTGTGCTGG; ACT2_qpcr_R: CAATTTCCCGCTCTGCTGTTG).

### Western-blotting

For western blot analysis, four-day-old seedling roots were collected, and ground in PBS buffer (pH7.4) containing protease inhibitors (Sigma, cOmplete, 11836170001). The extracts were mixed with 4× SDS loading buffer containing 5 mM DTT and boiled at 95°C for 5 min. Prior to immunoblotting, protein loading was adjusted based on 10% SDS-PAGE to ensure comparable loading among samples. mCherry-fusion proteins were detected using a rabbit anti-mCherry primary antibody (Sigma, SAB4702107) followed by a Sheep Anti-Rabbit IgG alkaline phosphatase (AP)-conjugated anti-rabbit secondary antibody (Rockland, 611-6502).

### Measurement of root hair length

The mature differentiation zone of 4-day-old (4 DAG) seedlings was imaged using an Olympus MVX10 microscope. For each root, the lengths of 10 fully elongated root hairs were measured in ImageJ and averaged. The number of seedlings used for statistical analysis is indicated in the corresponding figure legends.

### Use of artificial intelligence

D.J.C. used Nature Research Assistant Beta for shortening the abstract and identifying unclear technical details in the results section which were edited manually.

## Acknowledgments

We thank Edward Wagner, Daniel Durachko and Changhao Li for technical assistance. We also thank Hyung-Taeg Cho of Seoul National University for providing the GUS-GFP reporter vector. **Funding**: This work was supported by US Department of Energy, Office of Science, Basic Energy Sciences, under grant DE-FG02-84ER13179. N.K.H. was supported in part by a Graduate Research Fellowship from the US National Science Foundation (Grant No. DGE1255832). All data are available in the manuscript or the supplementary materials or posted in the Penn State Scholar Sphere repository DOI XXXXXXXXXX (# available upon publication)

