## Supplemental Figures for "Functional divergence of plant expansins as revealed by genetic complementation of root-hair tip growth in *Arabidopsis thaliana*"

**This PDF file includes:**

Figures S1 to S14
SI References

### Figures

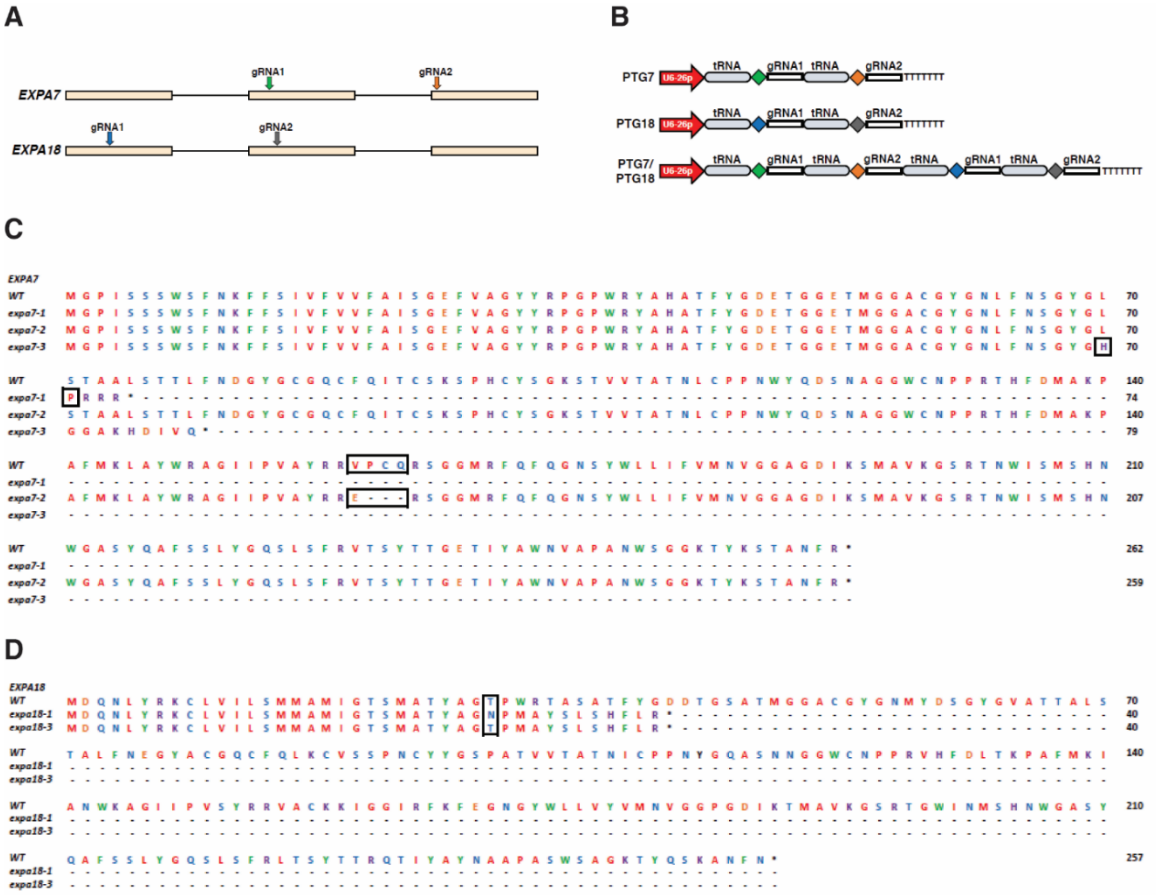

**Fig. S1. CRISPR/Cas9-mediated knockout of root hair–specific *EXPA7* and *EXPA18*.** (**A**) Schematic gene structures of *EXPA7* and *EXPA18* showing gRNA target sites (colored arrows). Exons are shown as tan boxes and introns as black lines. (**B**) Polycistronic tRNA-gRNAs (PTG) constructs targeting EXPA7-only (PTG7), EXPA18-only (PTG18), and both genes (PTG7/18). Target sequences indicated by arrows in (A) are represented as rhombi in matching colors. (**C** to **D**) Deduced amino acid sequences encoded by edited *EXPA7* (C) and *EXPA18* (D) alleles. Positions of sequence alterations are boxed in black. In *expa7-1*, a single guanine deletion (at 210 of *EXPA7* coding sequence) caused a reading frame (ORF)-shift, resulting in a truncated protein with 74 amino acid residues; *expa7-2* exhibited a 9-bp deletion from 479 to 487 of *EXPA7* coding sequence, losing three amino acid residues in the mature protein; *expa18-1* had a single adenine insertion between 88 and 89 of the *EXPA18* coding sequence, causing a truncated protein with 40 amino acid residues due to a frameshift; *expa7-3/expa18-3* had a 5-bp deletion from 209 to 213 of *EXPA7 coding sequence*, resulting in a truncated protein with 79 amino acid residues and a single cytosine insertion between 88 and 89 of EXPA18 coding sequencing, resulting in a truncated protein with 40 amino acid residues due to a frameshift.

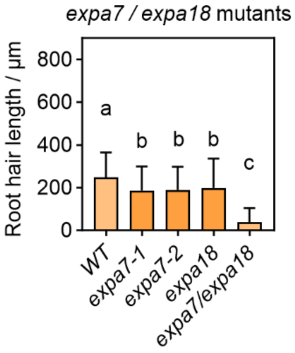


**Fig. S2.** Root hair length of single knock-out mutant in suboptimal growth conditions. Data are means ± SD (n=47, 54, 48, 59, 23; *, P<0.05; **, P<0.01; ****, P<0.0001; Student's t-test). Growth condition: ½ Murashige and Skoog (MS) medium supplemented with 1.5% sucrose and 0.6% Phytagel, pH 5.8 (1g/L MES), four-day growth under 16/8 hours light at 22/16°C.

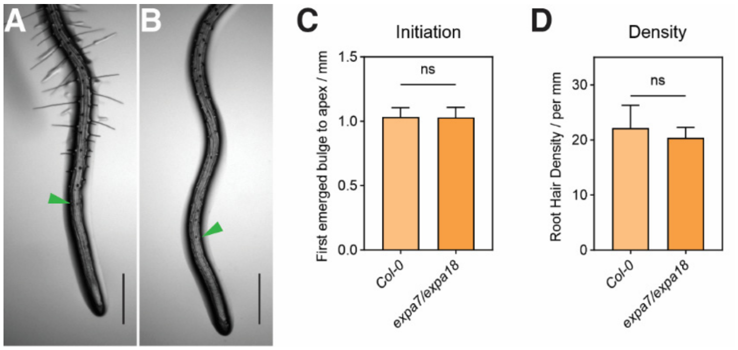


**Fig. S3. Knockout of *EXPA7* and *EXPA18* specifically impairs root hair elongation.** (**A** to **B**) Representative four-day-old root tips of Col-0 (A) and *expa7/expa18* (B); green arrowheads mark the first visible root hair bulges. Scale bars, 500 µm. (**C**) Distance from the root tip apex to the first visible root hair bulge; n = 13 (Col-0) and n = 11 (*expa7/expa18*). (**D**) The root hair (or root hair bulges) density per mm at the differentiation zone; means ± SD; n = 47 (Col-0) and n = 58 (*expa7/expa18*). Data are means ± SD; ns, not significant (P > 0.05; Student's t test).

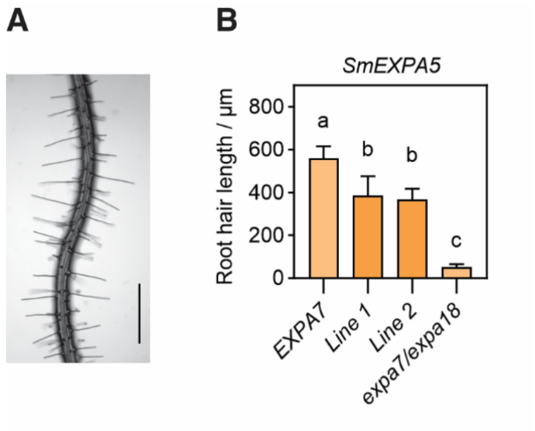


**Fig. S4. Restoration of root hair elongation in the *expa7/expa18* double mutant by *SmEXPA5* from *Selaginella moellendorffii*.** (**A**) Representative root hairs of *expa7/expa18* expressing *EXPA7pro::SmEXPA5*. Scale bar, 500 µm. (**B**) Root hair length in *EXPA7pro*::*SmEXPA5* lines 1 and 2 (n = 34 and 34, respectively), with *EXPA7pro*::*EXPA7* line 1 in the *expa7/expa18* background as a positive control (n = 30) and *expa7/expa18* as a negative control (n = 37). Data are means ± SD. One-way ANOVA followed by Tukey's HSD multiple comparison test; different letters denote statistically significant differences ($P < 0.0001$).

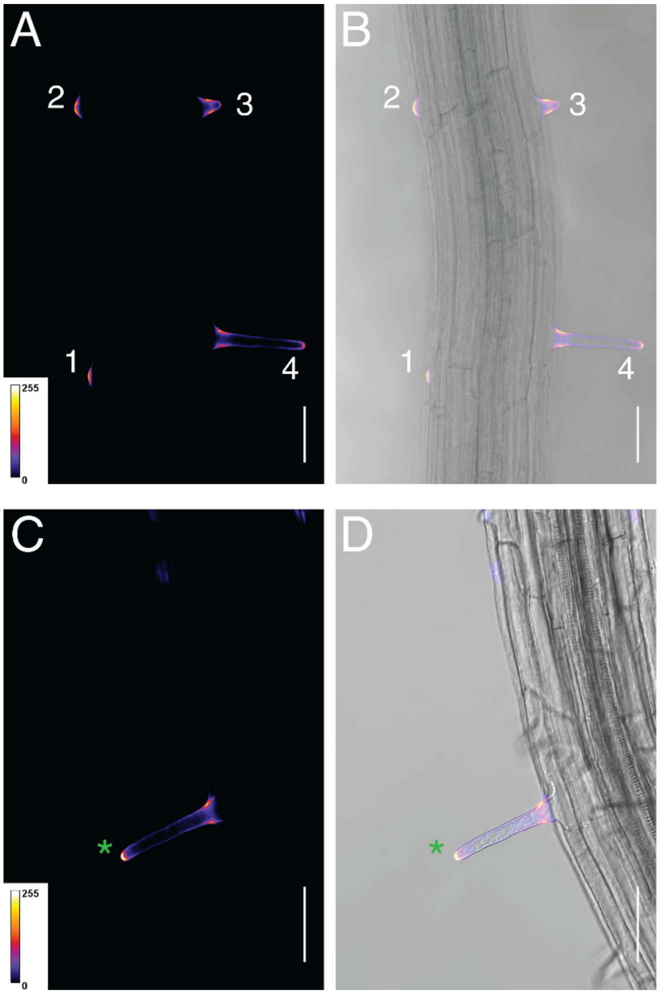


**Fig. S5. EXPA7-mCherry accumulation at the root hair bulge and tip apex.** (**A** and **B**) EXPA7-mCherry localization during early root hair development. Panel A shows the fluorescence channel, and panel B shows the merged fluorescence and bright-field image. Numbers indicate representative stages: bulging root hair (1), bulged root hair (2), transition stage (3), and tip growth stage (4). (**C** and **D**) EXPA7-mCherry localization in an elongated root hair. Panel C shows the fluorescence channel, and panel D shows the merged fluorescence and bright-field image. The green asterisk marks EXPA7-mCherry accumulation at the root hair tip. Fluorescence signals are displayed using the Fire LUT, in which pixel intensities are pseudocolored from low to high relative fluorescence signal, with darker colors indicating lower signal and warmer colors indicating higher signal; color bars indicate the displayed intensity range from 0 to 255.

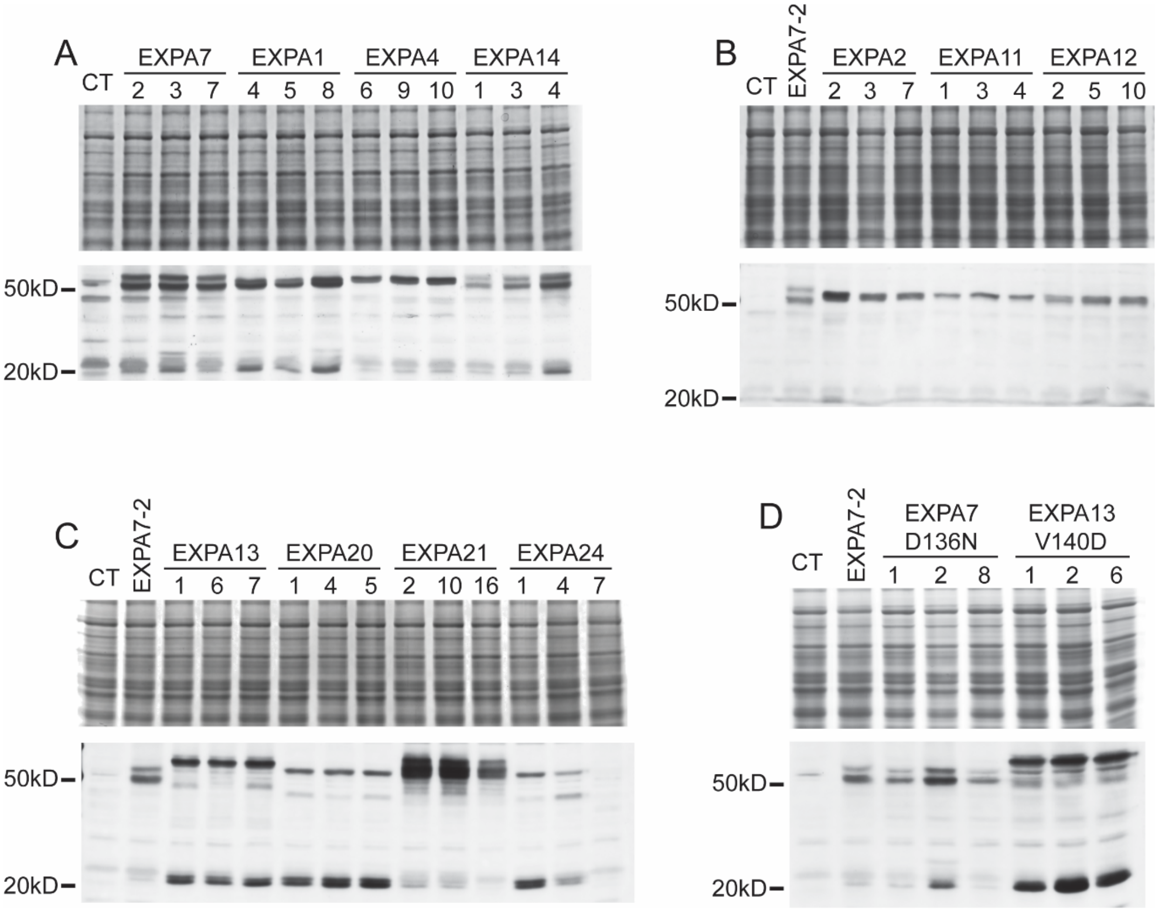


**Fig. S6. Protein detection of Chimeric mCherry fusions.** Total protein extracts from seedling roots expressing the indicated chimeric mCherry-fusion constructs were analyzed by SDS-PAGE and western blot. For each panel, the upper gel shows Coomassie Brilliant Blue staining, used to assess total protein loading, and the lower panel shows western blot detection of mCherry-fusion proteins by anti-mCherry antibody. CT indicates the control sample (expa7/expa18 mutant). Molecular mass markers are indicated on the left. (**A)** Detection of chimeric mCherry fusions derived from EXPA7, EXPA1, EXPA4, and EXPA14 constructs. (**B**) Detection of chimeric mCherry fusions derived from EXPA2, EXPA11, and EXPA12 constructs, with EXPA7-2 included as a reference sample. (**C**) Detection of chimeric mCherry fusions derived from EXPA13, EXPA20, EXPA21, and EXPA24 constructs, with EXPA7-2 included as a reference sample. (**D**) Detection of chimeric mCherry fusions carrying point mutations in EXPA7-D136N and EXPA13-V140D constructs, with EXPA7-2 included as a reference sample. Numbers above lanes indicate individual lines. Seedling age=four days after germination. Predicted molecular weight of mature EXPA proteins and mCherry: EXPA1, 27.0kD; EXPA2, 24.7kD, EXPA4, 25.8kD; EXPA7, 25.6kD; EXPA11, 24.8kD; EXPA12, 25.2kD; EXPA13, 26.9kD; EXPA20, 25.2kD; EXPA21, 24.6kD; EXPA24, 30.6kD; mCherry, 26.7KD.

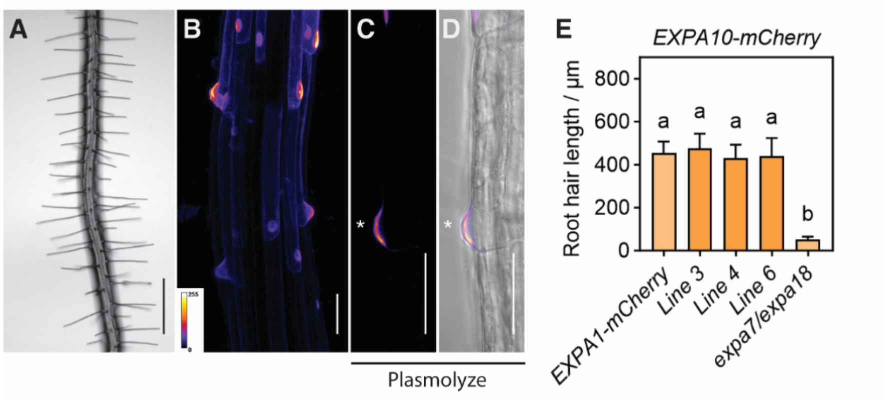


**Fig. S7. Root hair growth restoration and subcellular localization of EXPA10-mCherry in the *expa7/expa18* double mutant.** (**A** to **D**) Restoration of root hair growth and subcellular localization of EXPA10-mCherry in the *expa7/expa18* background. In (A to D), panels from left to right show root hairs; a maximum-intensity confocal projection of the root region spanning the transition from the elongation zone to the differentiation zone; the fluorescence channel; and merged fluorescence and bright-field channels of a plasmolyzed trichoblast. Asterisks mark the plasmolyzed cell wall at the RHID. Scale bars, 500 µm (A); 50 µm (B to D). Fluorescence signals are displayed using the Fire LUT. (**E**) Root hair length of independent lines expressing mCherry-fused EXPA10 (lines 3, 4, and 6; n = 34, 28, 42, respectively), with EXPA1-mCherry as a positive control (n = 50) and *expa7/expa18* as a negative control (n = 37). Data are presented as means ± SD. One-way ANOVA followed by Dunnett's multiple comparison test; different letters denote statistically significant differences ($P < 0.0001$).

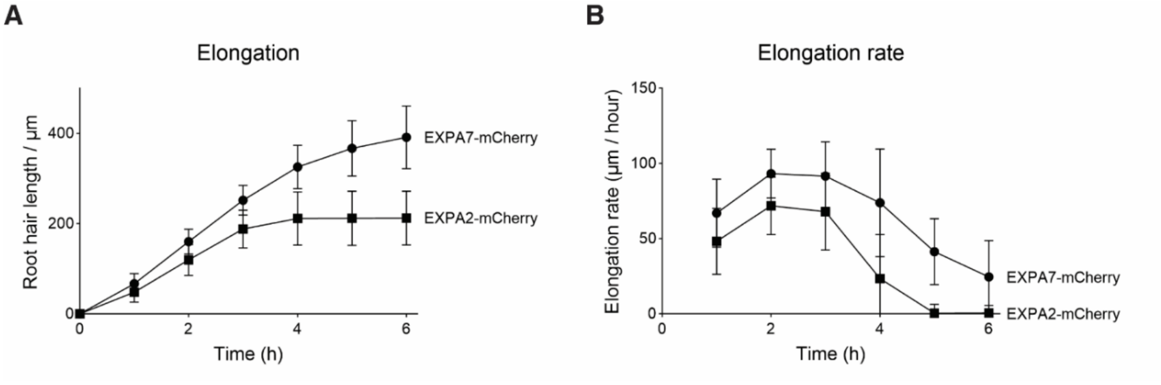


**Fig. S8. Root hair elongation in *expa7/expa18* lines complemented with EXPA7-mCherry or EXPA2-mCherry.** (**A**) Root hair length over time. The same individual root hairs from EXPA2-mCherry (squares) and EXPA7-mCherry (circles) were measured hourly throughout the observation period. (**B**) Elongation rate of the same root hairs, calculated from the change in length between consecutive time points.. Data are means ± SD; n = 15 (EXPA7-mCherry) and 20 (EXPA2-mCherry).

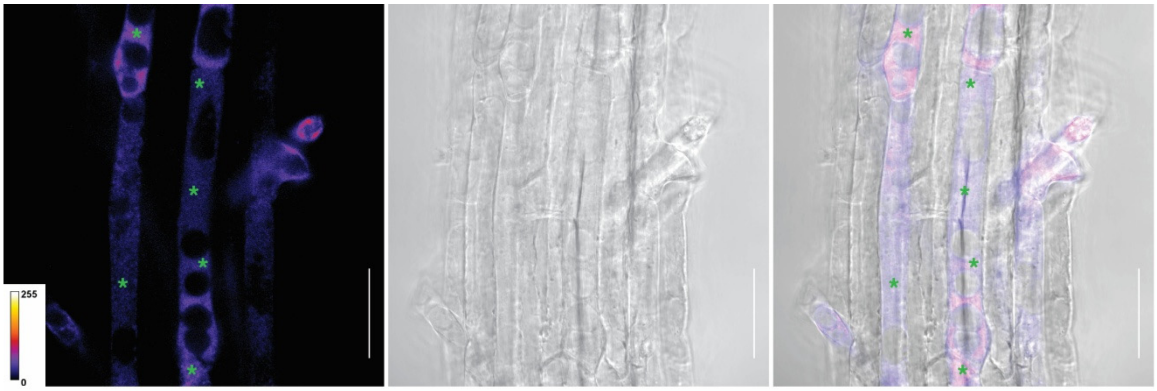


**Fig. S9. Subcellular localization of EXPA11-mCherry in plasmolyzed trichoblasts**. Four-day-old seedling roots expressing *EXPA7pro::EXPA11-mCherry* were plasmolyzed in 500 mM mannitol and imaged by confocal microscopy. The fluorescence channel (**A**), bright-field channel (**B**), and merged fluorescence and bright-field image (**C**) are shown. Green asterisks mark EXPA11-mCherry fluorescence in the apoplastic space between the cell wall and the retracted plasma membrane. Fluorescence signals are displayed using the Fire LUT. Scale bars, 50 μm.

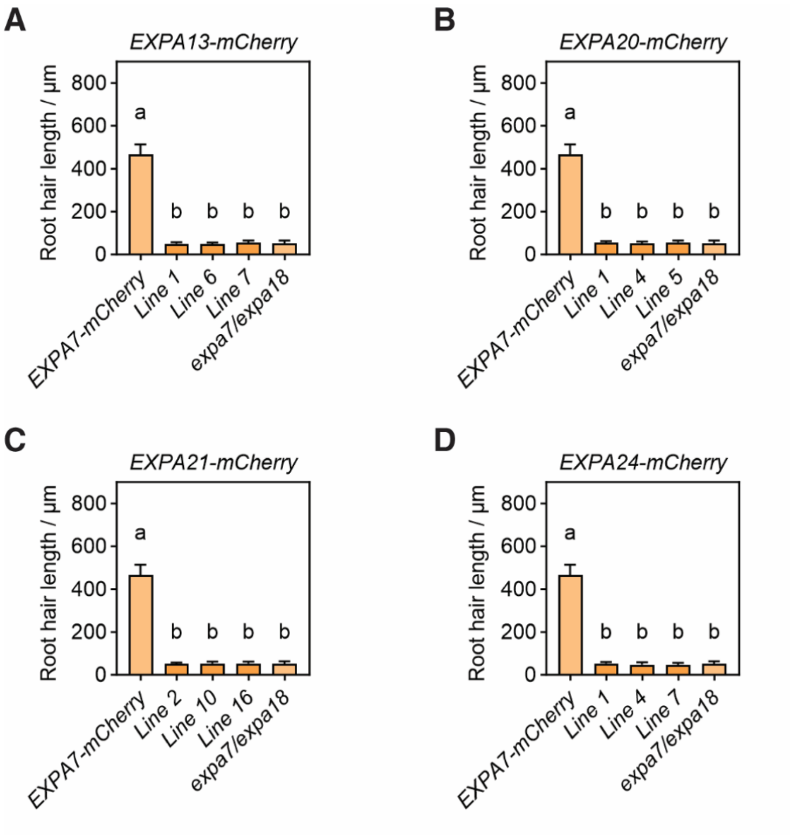


**Fig. S10. Lack of root hair growth restoration by mCherry-fused EXPA13, EXPA20, EXPA21, and EXPA24 in the *expa7/expa18* double mutant.** (**A** to **D**) Root hair length of independent lines expressing mCherry-fused EXPA13 (A, n = 47, 49, 44), EXPA20 (B, n = 30, 38, 29), EXPA21 (C, n = 51, 47, 40), and EXPA24 (D, n = 40, 40, 36); with EXPA7-mCherry line 1 as a positive control (n = 28) and *expa7/expa18* as a negative control (n = 37). Data are presented as means ± SD. One-way ANOVA followed by Tukey's HSD multiple comparison test; different letters denote statistically significant differences ($P < 0.0001$).

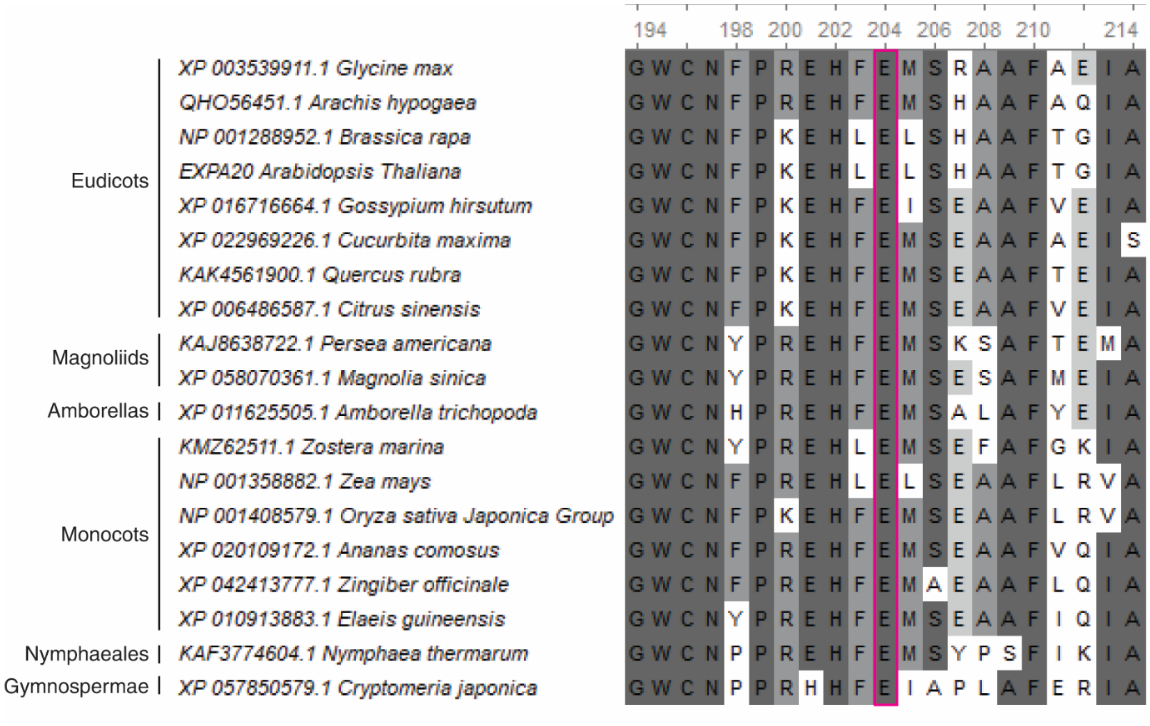

**Fig. S11. Sequence alignment of EXPA20 orthologs.** Amino acid sequence alignment (partial) of EXPA20 orthologs from representative plant species, grouped by major plant clades as indicated on the left**.** Residues corresponding to the conserved Asp (D) position in the HFD motif are replaced by Glu (E) in EXPA20 and its orthologs and are boxed in purple.

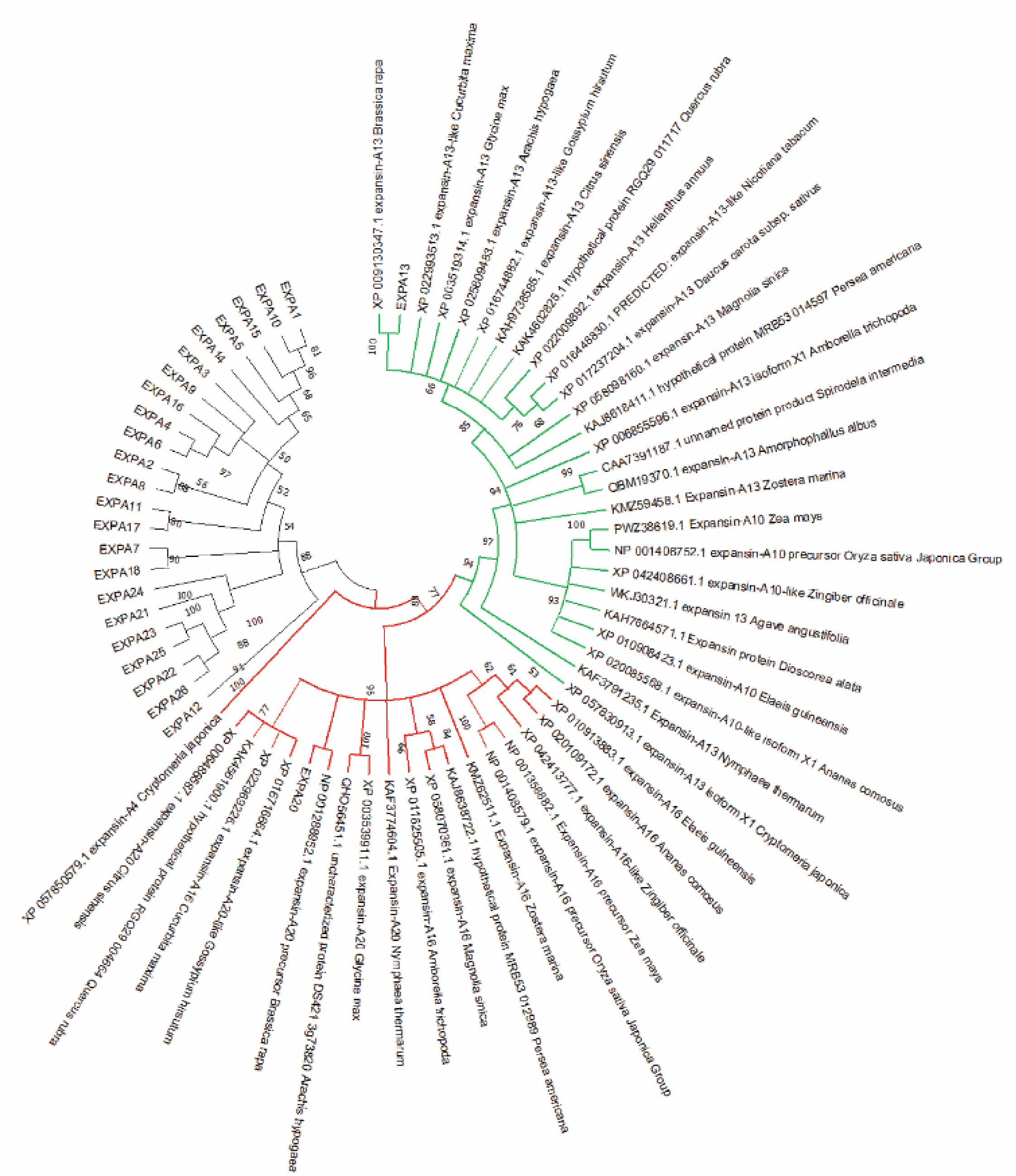

**Fig. S12. Unrooted neighbor-joining phylogenetic tree of Arabidopsis EXPAs and EXPA13/EXPA20 orthologs.** All Arabidopsis EXPA proteins together with EXPA13 and EXPA20 orthologs from gymnosperms, Amborellales, Nymphaeales, monocots, magnoliids, and eudicots (as in Fig. 4B and Fig. S8) were aligned and used to construct an unrooted neighbor-joining tree(1). Bootstrap support values from 10,000 replicates are shown at key nodes(2). Branches corresponding to EXPA13 orthologs are highlighted in green and those corresponding to EXPA20 orthologs in red. Evolutionary distances were computed using the Poisson correction method (3) and are expressed as the number of amino acid substitutions per site. Ambiguous positions were removed for each sequence pair (pairwise deletion), resulting in 413 positions in the final alignment of 50 protein sequences. The proportion of informative sites for each internal node is indicated next to the corresponding node. Phylogenetic analyses were performed in MEGA11(4).

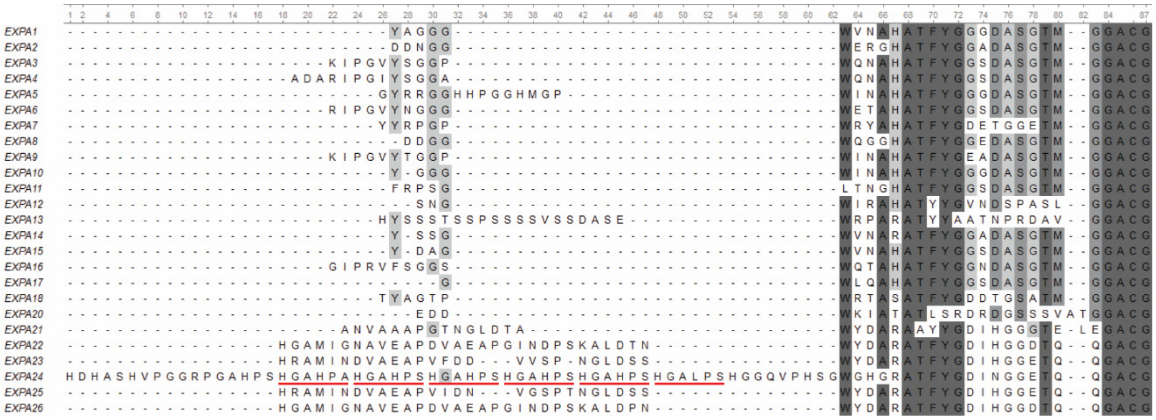

**Fig. S13. N-terminal region of the Arabidopsis EXPA mature protein alignment.** Multiple sequence alignment of Arabidopsis EXPA mature proteins after removal of the predicted signal peptides. The displayed region corresponds to the N terminus of the mature proteins. Red underlines mark the repetitive low-complexity region in EXPA24.

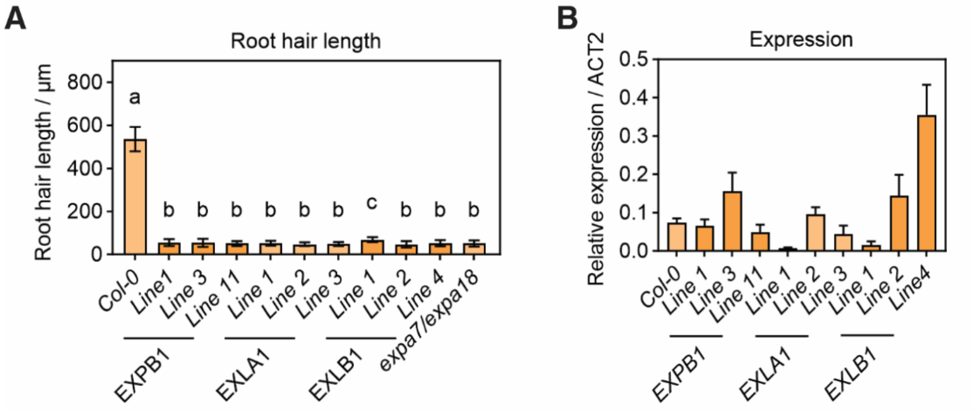


**Fig. S14. Failure of root hair growth complementation by EXPB1, EXLA1, and EXLB1 in the *expa7/expa18* double mutant.** (**A**) Root hair length in Col-0, *expa7/expa18*, and *expa7/expa18* lines expressing *EXPA7pro::EXPB1* (lines 1, 3, 11; n = 38, 34, 38), *EXPA7pro::EXLA1* (lines 1, 2, 3; n = 25, 27, 34), or *EXPA7pro::EXLB1* (lines 1, 2, 4; n = 59, 52, 26). None of these constructs restores root hair length to Col-0 levels; EXPA7pro::EXLB1 line 1 shows only a slight but significant increase over *expa7/expa18* (~17 µm; $P < 0.0001$). Data are presented as means ± SD. One-way ANOVA followed by Dunnett's multiple comparison test; different letters denote statistically significant differences ($P < 0.0001$). (**B**) Relative transcript levels determined by quantitative reverse-transcription PCR (qRT-PCR) for three independent lines of *EXPA7pro::EXPB1* (n = 4 per line), *EXPA7pro::EXLA1* (n = 4 per line), and *EXPA7pro::EXLB1* (n = 3 per line). Native *EXPA7* expression in Col-0 is shown for comparison; expression levels are normalized to *ACT2*.
